# U7 small nuclear RNA splice-switching therapeutics for STMN2 and UNC13A in Amyotrophic Lateral Sclerosis

**DOI:** 10.1101/2025.11.26.690143

**Authors:** Puja R. Mehta, Tomas Solomon, Sarah Pickles, Peter Harley, Michela Barioglio, Christoph Schweingruber, Alessandro Marrero-Gagliardi, Yujing Gao, Francesca Mattedi, Simone Barattucci, Lilian Tsai-Wei Lin, Eugeni Ryadnov, Matteo Zanovello, Alexander J. Cammack, Adrian M. Isaacs, Juan Burrone, Christopher E. Shaw, Matthew J. Keuss, Leonard Petrucelli, Pietro Fratta, Marc-David Ruepp

## Abstract

TDP-43 nuclear depletion in amyotrophic lateral sclerosis (ALS) causes de-repression of cryptic exons (CEs) in multiple transcripts, including *UNC13A* and *STMN2*, disrupting synaptic transmission and neurite outgrowth. We developed a therapeutic U7 snRNA (tU7) approach that suppresses TDP-43-dependent mis-splicing, restores target gene expression, rescues neuronal functions in human iPSC-derived neurons, and shows target engagement *in vivo*, positioning tU7-mediated splicing correction as a promising therapeutic strategy for ALS.

## Introduction

Cytoplasmic mislocalisation accompanied by nuclear depletion of the RNA-binding protein, TDP-43, represents the most common pathological feature of amyotrophic lateral sclerosis (ALS), occurring in the vast majority (>97%) of cases ^1^. Nuclear loss of TDP-43 disrupts RNA processing, producing widespread splicing defects including the de-repression of intronic sequences that are aberrantly incorporated in mature RNA – termed cryptic exons (CEs) ^2^. Evidence from multiple groups indicates that these CEs are not just disease epiphenomena but active contributors to disease development and progression ^3,4^. Inclusion of these CEs often results in loss of functional mRNAs and their encoded proteins ^2,5^. UNC13A and STMN2 are two proteins that are depleted via CE-mediated mis-splicing and have critical neuronal functions which are relevant to disease pathogenesis: UNC13A is a synaptic protein required for neurotransmitter release and transmission of nerve impulses across the synapse ^3,4,6^, and STMN2 is a microtubule-associated axonal protein crucial for neurite outgrowth ^7,8^ and whose loss results in a slowly progressive, motor-selective neuropathy with functional deficits and neuromuscular junction denervation ^9^.

These discoveries have inspired the design of antisense oligonucleotides (ASOs) to correct these CE splicing events individually ^6,10^. Although these strategies may slow disease progression, the occurrence of numerous CEs in human disease characterised by TDP-43 proteinopathy suggests that correcting multiple mis-splicing events would be beneficial. Targeting multiple CEs simultaneously with ASOs, however, may be constrained by cumulative oligonucleotide-associated toxicity. An alternative approach to modulate splicing is the use of vectorised modified U7 small nuclear RNAs (U7smOPT) ^11^, which are currently under evaluation in a clinical trial for Duchenne muscular dystrophy ^12^. Owing to their compact size, multiple therapeutic U7 (tU7) cassettes can be incorporated into a single adeno-associated viral (AAV) vector ^13,14^, thereby enabling the targeting of multiple splicing events and providing a sustained therapeutic effect. We therefore investigated the potential of U7 snRNAs to repress CE splicing as a therapeutic modality for ALS with TDP-43 proteinopathy.

## Results and discussion

### tU7s rescue UNC13A and STMN2 cryptic exon inclusion

We generated a series of U7 smOPT snRNAs ^11^ targeting the TDP-43 binding regions (TDPbr), expected to be accessible upon nuclear TDP-43 loss, to restore CE repression in *UNC13A* and *STMN2*. In addition to the targeting antisense sequences, these snRNAs contain a short non-complementary RNA tail harbouring two canonical motifs designed to recruit hnRNPA1 to substitute for TDP-43’s suppressive function (**Fig. 1a**). We initially generated eight such bifunctional TDPbr-targeting U7 smOPT snRNAs for *UNC13A*, and seven for *STMN2*, along with ones targeting the proximal 3’-splice site in *UNC13A* and a putative exonic splicing enhancer in *STMN2*. We assessed their efficacy using *UNC13A* and *STMN2* minigenes in TDP-43-depleted 293T cells. These experiments demonstrated the efficacy of the TDPbr-targeting approach (**Fig. S1a-d**). In addition, we investigated whether multiple CEs can be targeted in parallel and engineered a construct containing three tU7 cassettes directed against CEs in *UNC13A* and *STMN2*, as well as against a CE in *INSR* ^*2*^. This approach yielded a level of CE suppression in *UNC13A* and *STMN2* comparable to that achieved by targeting each exon individually, demonstrating that multiple tU7 cassettes can be combined to enable simultaneous correction of CE splicing (**Fig. S1e-g**).

**Figure 1.**
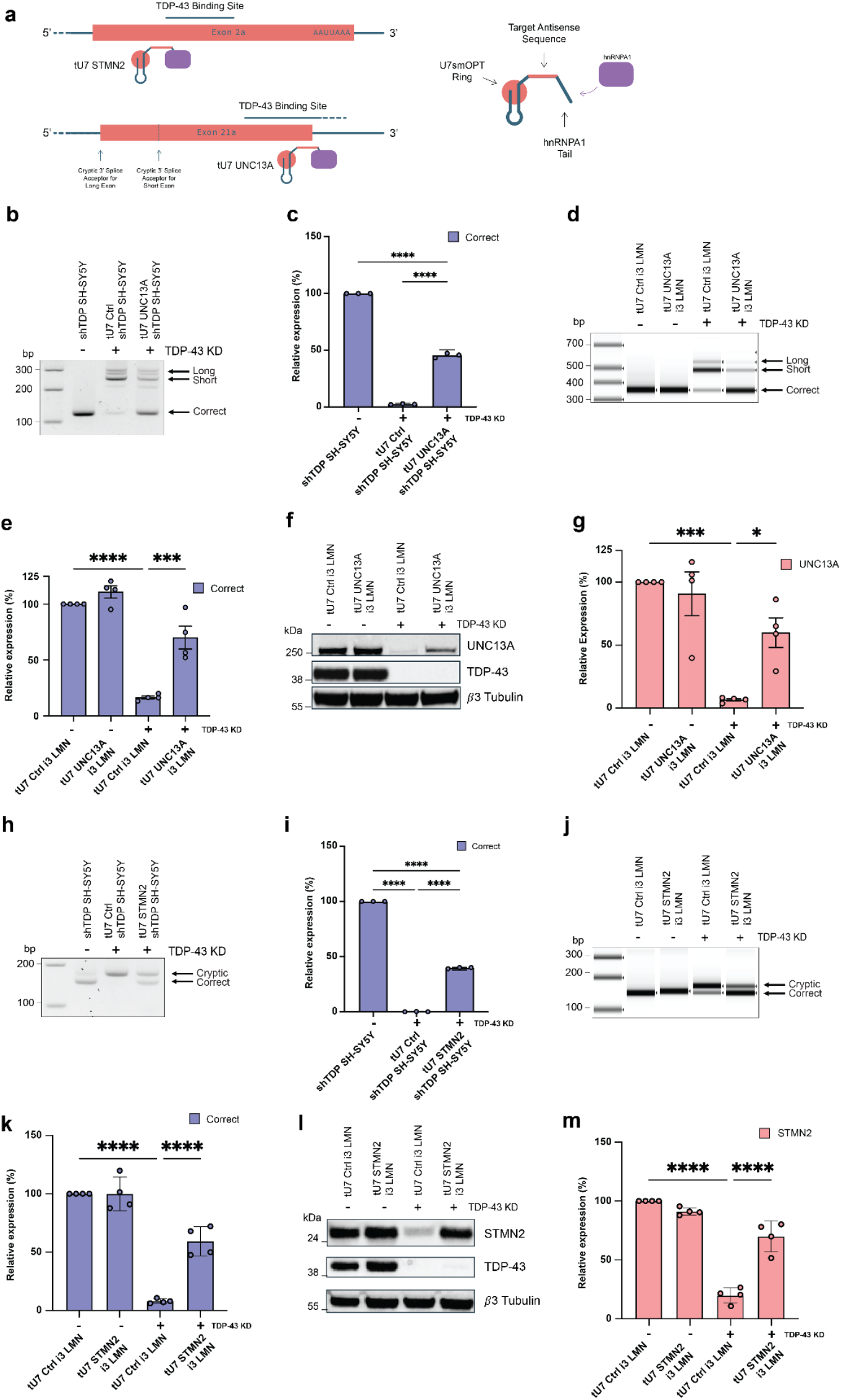
tU7s rescue *UNC13A* and *STMN2* mis-splicing in SH-SY5Y and i3-LMNs and restore protein levels. **a**, Schematic representation of tU7s and respective binding regions. **b, h**, Representative image of RT-PCR products showing *UNC13A* (**b**) and *STMN2* (**h**) correct and cryptic mature mRNA levels in SH-SY5Y cells with non-targeting tU7 Ctrl and targeting tU7 constructs following shRNA mediated TDP-43 knockdown (KD). The respective tU7 constructs are able to partially rescue mis-splicing. **c, i**, Quantification (mean ± SEM) of RT-qPCR levels of *UNC13A* (**c**) and *STMN2* (**i**) correctly spliced mature mRNA relative to tU7 Ctrl without TDP-43 KD in the SH-SY5Y line (*n* = 3 biological replicates). **d, j**, Representative image of RT-PCR products showing *UNC13A* (**d**) and *STMN2* (**j**) correct and cryptic mature mRNA levels in i3-LMNs with non-targeting tU7 Ctrl and targeting tU7 constructs following CRISPRi-mediated TDP-43 KD. The respective tU7 constructs are able to partially rescue mis-splicing. **e, k**, Quantification (mean ± SEM) of RT-qPCR levels of *UNC13A* (**e**) and *STMN2* (**k**) correctly spliced mature mRNA relative to tU7 Ctrl without TDP-43 KD in the i3 LMN line (*n* = 4 biological replicates). **f, g, l, m**, Western blot demonstrating UNC13A (**f**) and STMN2 (**l**) protein loss in CRISPRi-mediated TDP-43 KD and rescue using the tU7s (*n* = 4 biological replicates). Quantification (mean ± SEM) of UNC13A (**g**) and STMN2 (**m**) protein levels relative to tU7 Ctrl without TDP-43 KD, showing the rescue of UNC13A and STMN2 protein after treatment with the respective tU7 (*n* = 4 biological replicates). Statistical significance for **c, e, i, k, g, m** was determined by one-way ANOVAs with Tukey multiple comparison (*p < 0.05; **p < 0.01; ***p < 0.001; ****p < 0.0001; ns = non-significant).

### tU7s rescue endogenous cryptic splicing and protein levels of UNC13A and STMN2 in i3-LMNs

In order to assess efficacy on the endogenous RNAs, which are only expressed in neuronal cells, the best performing TDPbr-targeting tU7s for *UNC13A* and *STMN2* were then taken forward individually for validation in neuronal models. We generated lentiviral vectors harbouring either an *UNC13A, STMN2*, or non-targeting control (Ctrl) tU7 expression cassette alongside an EF1a-driven mCherry-T2A-BSD cDNA. After transduction of SH-SY5Y neuroblastoma cells harbouring a doxycycline-inducible TDP-43 shRNA cassette, the induction of TDP-43 knockdown led to an increase in CE-containing *UNC13A* mRNA at the expense of correctly spliced *UNC13A* in the presence of the non-targeting tU7 Ctrl. This defect was rescued by the expression of the tU7 *UNC13A* (**Fig. 1b**). In line with this observation, RT-qPCR demonstrated that TDP-43 depletion in the presence of the Ctrl tU7 reduced relative *UNC13A* mRNA levels to 2.4%, whereas tU7 UNC13A expression restored transcript levels to ~50% (**Fig. 1c**). The same effect was observed in human iPSC-derived lower motor neurons (i3-LMNs) after CRISPRi-mediated TDP-43 depletion (**Fig. 1d-e**), where tU7 UNC13A suppressed CE inclusion and restored correctly spliced *UNC13A* mRNA to ~70%. In line with this correction, UNC13A protein levels in i3-LMNs were restored to ~60% (**Fig. 1f-g**). Similarly, *STMN2* correction was achieved in SH-SY5Y and i3-LMNs as observed by RT-PCR (**Fig. 1h, j**) with ~40% and ~60% rescue of correct *STMN2* mRNA, respectively (**Fig. 1i, k**). In line with this correction, this resulted in the restoration of STMN2 protein levels in i3-LMNs to ~70% (**Fig. 1l-m**). Given that restoration to >25% of STMN2 levels rescues axonal phenotypes ^10^ and >30% of UNC13A levels rescue synaptic deficits ^6^, we concluded that these constructs achieve therapeutically relevant restoration of UNC13A and STMN2 levels.

### UNC13A tU7s rescue synaptic function

We thus advanced towards assessing whether tU7s achieve phenotypic rescue. To assess restoration of synaptic deficits induced by UNC13A loss, we used homozygously Halo-tagged TDP-43 iPSCs (Halo-i3N) lines in which TDP-43 can be depleted through HaloPROTAC treatment ^6^. Cells were transduced with tU7 Ctrl, tU7 UNC13A, and tU7 STMN2 lentiviral vectors. Non-transduced and tU7-transduced Halo-TDP-43 iPSCs were differentiated to cortical-like i3Neurons and assessed in the presence or absence of TDP-43 depletion. Semi-quantitative RT-PCR (**Fig. 2a**) demonstrated a reduction of correctly spliced *UNC13A* mRNA at the expense of cryptic *UNC13A* mRNA, accompanied by a reduction in total *UNC13A* mRNA levels, as measured by RT-qPCR (**Fig. S2a**). Quantitative assessment by RT-qPCR (**Fig. 2b**) demonstrated a reduction of correctly spliced *UNC13A* mRNA to 4% in TDP-43-depleted neurons, which was restored to >50% in tU7 UNC13A-transduced cells relative to non-HaloPROTRAC treated cells. This rescue on the RNA level translated to a restoration of UNC13A protein levels from 1% to 64% of levels observed in the presence of TDP-43 (**Fig. 2c-d**), in turn leading to a restoration of UNC13A staining at synapsin-positive presynaptic terminals (**Fig. 2e-f**), thus re-instating a key molecular constituent required for synaptic function. More importantly, we found that impaired synaptic vesicle release, indicated by a reduction in the frequency of miniature excitatory postsynaptic currents (mEPSCs) in the absence of TDP-43, could be rescued by UNC13A tU7s (**Fig. 2g-h**).

**Figure 2.**
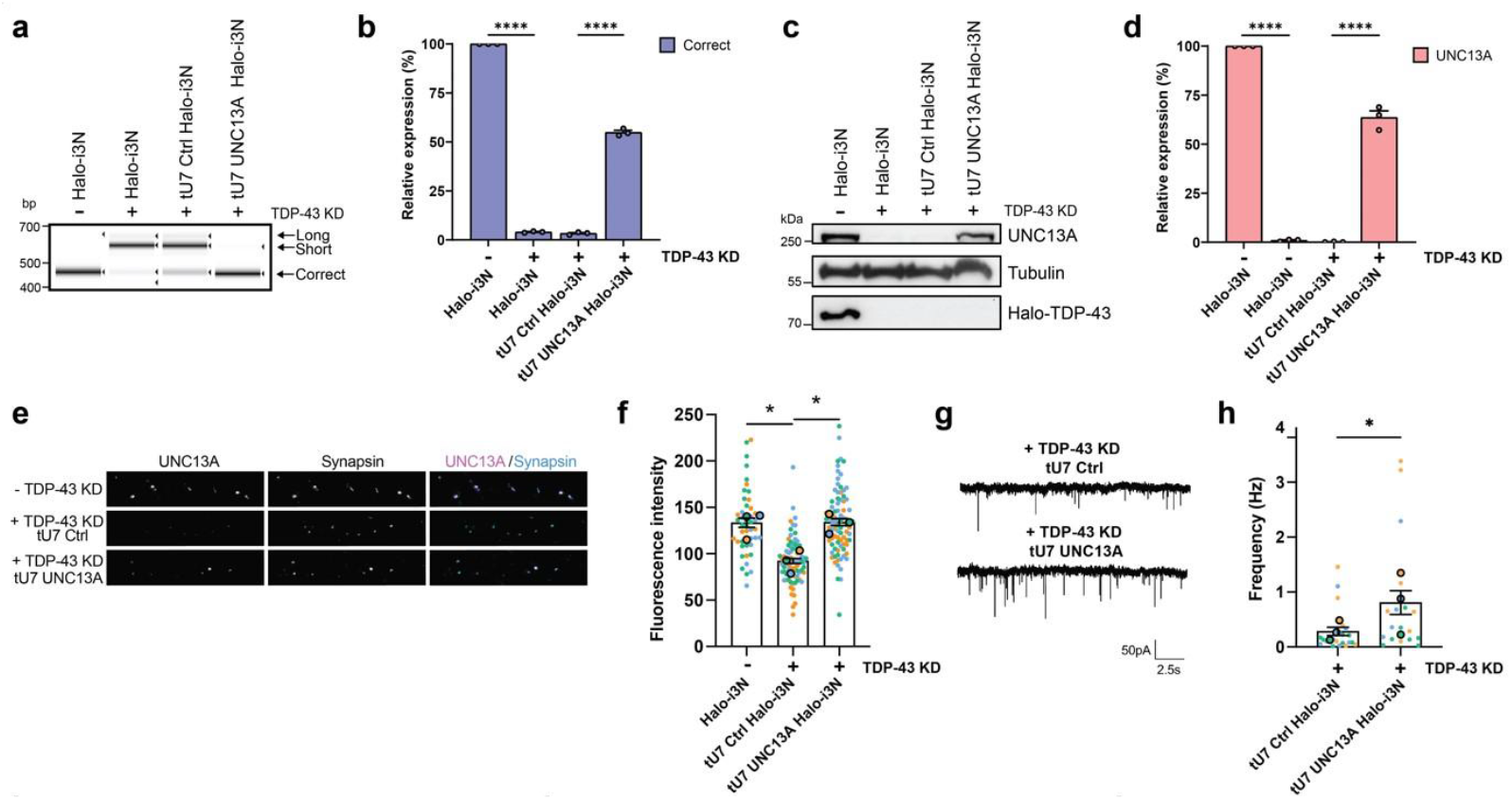
UNC13A tU7 rescues cellular mis-splicing phenotypes in TDP-43-depleted neurons. **a**, Representative image of RT-PCR products shows targeted tU7 inhibition of *UNC13A* cryptic exon inclusion compared to a non-targeting tU7 Ctrl, after degron-mediated TDP-43 KD in human iPSC-derived cortical-like i3Neurons (*n* = 3 biological replicates). **b**, Quantification (mean ± SEM) of RT-qPCR levels of correctly spliced *UNC13A* transcripts shows successful rescue of correctly spliced transcripts after tU7 treatment (*n* = 3 biological replicates). **c**,**d**, Representative western blot shows levels of UNC13A protein after targeted tU7 treatment compared to the tU7 Ctrl in i3Neurons after TDP-43 KD (*n* = 3 biological replicates). **d** Quantification (mean ± SEM) of protein levels normalised to tubulin shows successful rescue of UNC13A after tU7 treatment. **e**, Representative immunofluorescence labelling of synapsin and UNC13A at pre-synaptic terminals in 4-week-old i3Neurons grown on rat astrocytes following 2 weeks of TDP-43 KD. Scale bar = 10 μm. **f**, Quantification (mean ± SEM) of UNC13A intensity at synapsin-positive terminals shows tU7 UNC13A rescue of UNC13A at the pre-synaptic terminal. No TDP-43 KD *N*=71, TDP-43 KD + tU7 Ctrl *N*=71, TDP-43 KD + tU7 UNC13A *N*=66 synapses from *n* = 3 experiments. Each dot represents data from one synapse. Different colours represent independent differentiations. **g**, Representative mEPSC traces from tU7 Ctrl and tU7 UNC13A-expressing i3Neurons. **h**, Quantification (mean ± SEM) of mEPSC frequency from TDP-43 KD + tU7 Ctrl *N*=17 and TDP-43 KD + tU7 UNC13A *N*=22 i3Neurons pooled from *n* = 3 experiments shows tU7 UNC13A rescue of mEPSC frequency. Statistical significance for **b** and **d** was determined by one-way ANOVA followed by Tukey’s multiple comparisons test. Statistical significance for **f** was determined by one-way ANOVA followed by Dunnett’s multiple comparisons test, and **h** by a paired *t*-test (\**p* < 0.05; \*\**p* < 0.01; \*\*\**p* < 0.001; \*\*\*\**p* < 0.0001; ns = non-significant).

### STMN2 tU7s rescue neurite outgrowth deficits

We then assessed STMN2 tU7 in i3Neurons. The HaloTag intrinsically affected TDP-43-mediated *STMN2* CE repression, as observed by RT-PCR (**Fig. 3a**). Quantitative assessment by RT-qPCR demonstrated a ~50% reduction of correctly spliced and total *STMN2* mRNA in the presence of Halo-tagged TDP-43, which tU7 STMN2 restored to near wildtype levels. Upon TDP-43 depletion, correctly spliced *STMN2* levels were essentially absent, but were rescued to ~80% after tU7 STMN2 transduction (**Fig. 3b, Fig. S2b-c**). This rescue on the RNA level translated to a restoration of STMN2 protein from 3% to 70% of wildtype levels (**Fig. 3c-d**). Critically, we found that tU7-mediated restoration of STMN2 protein levels also rescues TDP-43 depletion-mediated neurite outgrowth impairment (**Fig. 3e-f**).

**Figure 3.**
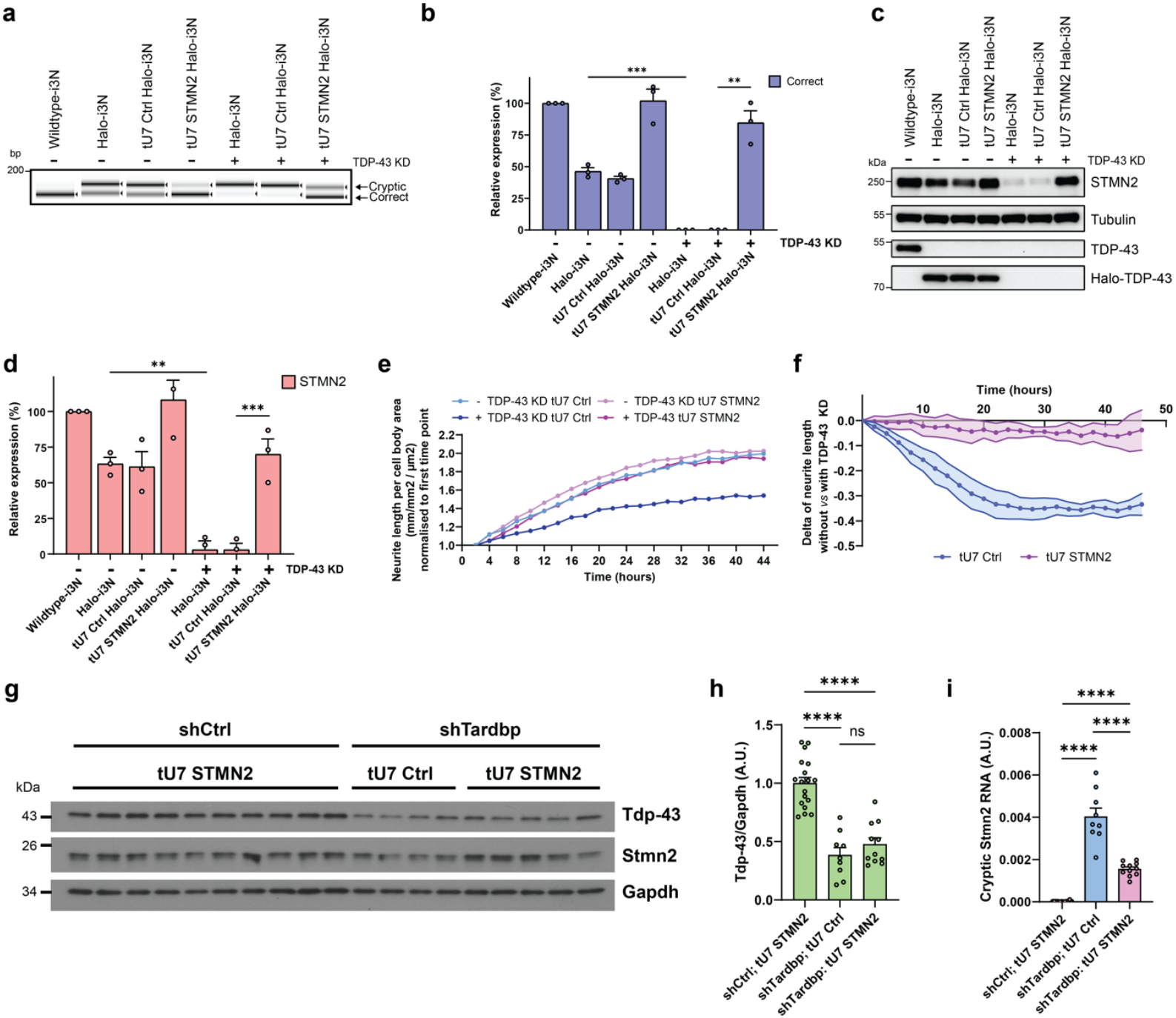
STMN2 tU7 rescues cellular mis-splicing phenotypes in TDP-43-depleted neurons and shows *in vivo* target engagement. **a**, Representative image of RT-PCR products shows targeted tU7 inhibition of *STMN2* cryptic exon inclusion compared to a non-targeting tU7 Ctrl, after degron-mediated TDP-43 KD in human iPSC-derived cortical-like i3Neurons (*n* = 3 biological replicates). **b**, Quantification (mean ± SEM) of RT-qPCR levels of correctly spliced and *STMN2* transcripts shows successful rescue of correctly spliced transcripts after tU7 treatment (*n* = 3 biological replicates). **c**,**d**, Representative western blot shows levels of STMN2 protein after targeted tU7 treatment compared to the tU7 Ctrl in i3Neurons after TDP-43 KD (*n* = 3 biological replicates). **d**, Quantification (mean ± SEM) of protein levels normalised to tubulin shows successful rescue of STMN2 after tU7 treatment. **e**, Representative graph of neurite outgrowth (plotted as neurite length per total cell body area normalised to the first time point) over time in tU7 STMN2-expressing and tU7 Ctrl-expressing i3Neurons without and with TDP-43 KD. **f**, Quantification (mean ± SEM) at each time point of the delta values between neurite outgrowth (neurite length per cell body area, normalised to the first time point) without *vs* with TDP-43 KD degron treatment. This was plotted for tU7 Ctrl (blue) and tU7 STMN2 (pink)-expressing i3Neurons, showing a greater neurite outgrowth impairment in tU7 Ctrl-expressing i3Neurons after TDP-43 KD compared to tU7 STMN2-expressing i3Neurons after TDP-43 KD (n = 5 biological replicates). **g-i**, RNA and protein were measured in the forebrains of 1.5-month-old humanised Stmn2^+/+^ mice ICV injected with AAV expressing shRNA and tU7 snRNA, *shCtrl;* tU7 STMN2 (*n* = 18-21 biological replicates), *shTardbp;* tU7 Ctrl (*n* = 9 biological replicates), and *shTardbp;* tU7 STMN2 (*n* = 11 biological replicates). Representative western blot **(g)** and quantification (mean ± SEM) **(h)** of Tdp-43 protein normalised to Gapdh levels, reveal a reduction of Tdp-43 protein in *shTardbp*-treated mice. **i**, Quantification (mean ± SEM) of cryptic *Stmn2* RNA, normalised to endogenous controls *Rplp0* and *Gapdh*, as measured by RT-qPCR, reveal a significant reduction in *shTardbp*; tU7 STMN2-treated mice. Statistical significance for **b**,**d**,**h**, and **i** was determined by one-way ANOVA followed by Tukey’s multiple comparisons test (\**p* < 0.05; \*\**p* < 0.01; \*\*\**p* < 0.001; \*\*\*\**p* < 0.0001; ns = non-significant).

### tU7s are effective in correcting cryptic splicing in vivo

Having demonstrated successful rescue of splicing, protein levels, and functional phenotypes *in vitro*, we next evaluated target engagement *in vivo*. Given that *UNC13A* and *STMN2* cryptic exons are not conserved between humans and mice, we utilised a humanised *Stmn2* mouse model (Stmn2^em6(STMN2)^) harbouring a homozygous knock-in of the human *STMN2* cryptic exon (exon 2a) ^10^. We performed P0 intracerebroventricular (ICV) injections with AAV vectors (AAV-PHP.eB) containing a short hairpin RNA (shRNA) expression cassette to deplete Tdp-43 (shTardbp) combined with either tU7 Ctrl or tU7 STMN2 expression cassettes, alongside a vector containing a non-targeting shRNA (shCtrl) and tU7 STMN2 cassette. RNA and protein were isolated from forebrains at 1.5 months post-injection. Assessment by western blot demonstrated a reduction in Tdp-43 protein levels to 37% (**Fig 3g-h**) and *Tardbp* mRNA to 32% (**Fig. S2d**) of shCtrl-treated samples. The reduction of Tdp-43 was accompanied by a significant increase in CE-containing *Sort1* and alternatively spliced *Tsn* mRNAs ^15,16^ (**Fig. S2e-f**) alongside a non-significant trend towards lower *Stmn2* mRNA and protein levels (**Fig. S2g-h**). Reduction of Tdp-43 resulted in a marked emergence of cryptic *Stmn2*-containing RNA (**Fig. 3i**), and this was significantly reduced in samples treated with the tU7 STMN2-containing *Tardbp* shRNA vector, demonstrating the effectiveness of tU7s to correct TDP-43-mediated cryptic splicing *in vivo*.

In conclusion, we demonstrate that tU7 snRNA-mediated cryptic splicing correction is a promising therapeutic option for ALS and potentially other central nervous system TDP-43 proteinopathies, such as frontotemporal dementia with TDP-43 pathology, Alzheimer’s disease, and limbic-associated TDP-43 encephalopathy (LATE) ^17^. This approach allows for the targeting of two different tU7 snRNA targets with long-term benefit *in vivo* ^18^ and holds promise for simultaneous targeting of multiple CE events with the potential to provide sustained benefit for the majority of people with ALS, for which there are currently no effective disease-modifying treatments available.

## Acknowledgements

This research was funded by the support from the UK Dementia Research Institute (award numbers DRI-TAP2021-11 to PF, CES and MDR, DRI-LTA2023/4 to PF and MDR, UK DRI-1203 to AMI and DRI-6204 to MDR) through UK DRI Ltd, principally funded by the UK Medical Research Council. PF is supported by UK Medical Research Council and MNDA Senior Clinical Fellowship and a Lady Edith Wolfson Fellowship (MR/M008606/1 and MR/S006508/1). PRM was supported by a Wellcome Trust Clinical Training Fellowship (102186/B/13/Z) and the MND Association (Biomedical Project Grant award 893-791 to PRM). SP is supported by an Alzheimer’s Association Research Fellowship (AARF-1308004) and an Innovation of Ageing Award from Mayo Clinic’s Center for Clinical and Translational Science and the Robert and Arlene Kogod Center on Ageing. PF is supported by the MRC (MR/W005190/1) AJC was supported by a Live Like Lou Foundation postdoctoral fellowship. JB was supported by the Wellcome Trust (215508/Z/19/Z), the UK Medical Research Council (MR/Z505353/1, to JB and PF) and the MND Association (2457-797 to JB and PF). LP is supported by the National Institutes of Health (U54NS123743) and the Cure Alzheimer’s Fund. We thank Daniel Schuemperli fur valuable discussions, Micheal Ward and Christopher Grunseich for the iPSC lines and Alessandro Bertero for sharing the pAAV-Puro_siKD plasmid.

## Contributions

Initial concept CES, PF, and MDR. PRM, CES, PF, and MDR secured funding. PF and MDR oversaw project administration. JB, AMI, MJK, LP, PF, and MDR provided supervision. Methodology and further study design was developed by PRM, TS, SP, PH, MJK, LP, PF, and MDR. Investigations were performed by PRM, TS, SP, PH, MB, CS, YG, FM, SB, MZ, AJC, LT-WL, and ER. Resources were provided by PRM, TS, SP, CS, AMG, AMI, MJK, JB, and LP. PRM and TS designed and prepared visualisations. The original draft was written by PRM, TS, PF, and MDR, and all authors contributed to the review and editing of the manuscript.

## Ethics declarations Competing interests

PF, CES, MDR have filed patent applications relating to the use of modified U7 snRNAs for the correction of TDP-43-regulated cryptic exons. PF is cofounder of Trace Neuroscience. PF and MK have filed patents related to the use of UNC13A as therapeutic target.

## Material and methods

### Design of therapeutic U7s (tU7s)

A series of bifunctional tU7 snRNAs were designed to target either the 3’-splice site (also named the splice acceptor site), 5’-splice site (also named the splice donor site), TDP-43 binding regions (identified using iCLIP data; ^19^), or putative exon splicing enhancer (ESE) regions (identified using ESE finder 3.0) for target genes. Details regarding antisense sequences and bifunctional tails are listed for all tU7 constructs in **Table S1** and **Figure S1**. Bifunctional tU7s contain an additional tail sequence for recruitment of a splicing repressor protein to promote cryptic exon-skipping (hnRNPA1).

The U7 SmOPT expression cassettes for testing in 293T cells were ordered as gene synthesis in pMK with a CMV-driven blasticidin resistance (GeneArt, Life Technologies). To generate the constructs targeting cryptic exons, these constructs were digested with StuI and HindIII (New England Biolabs). DNA strings with 15 bp overhangs upstream and downstream of the StuI, HindIII cleavage sites containing the U7 SmOPT sequence with the antisense sequence were designed as described above and cloned into the StuI and HindIII digested U7SmOPT plasmid using InFusion Snap Assembly EcoDry Master Mix (Takara) according to the manufacturer’s instructions.

### UNC13A and STMN2 Minigenes

The UNC13A minigene is described in^3^. The STMN2 minigene was generated by gene synthesis (GeneArt, Life Technologies) and assembled to include: (i) exon 1 with 300 bp of downstream intron 1; (ii) the cryptic exon 2a flanked by 300 bp of upstream intron 1 and 200 bp of downstream intron 1; (iii) exon 2 with 200 bp of upstream and 200 bp of downstream intronic sequence; and (iv) exon 3 with 200 bp of upstream intronic sequence. This fragment was cloned between the BamHI and XhoI sites of pcDNA3.1(+).

### Inducible TDP-43 knockdown 293T cells

293T cells were cultured in DMEM/F12 medium (Gibco) with 10% tetracycline-free FBS and 1% Penicillin/Streptomycin. Inducible 293T TDP-43 knockdown cells were generated by transfecting 80% confluent cells in a well of a 6-well plate with AAVS1-SA-puro-EF1-hspCas9 (System Biosciences) targeting the AAVS1 locus (ggggccactagggacaggat) and pAAVS1-puro 2x TDP-43 shRNA in a 1:3 ratio. The pAAVS1-puro 2x TDP-43 shRNA plasmid was generated by cloning a gene-synthetised fragment containing two Tet-operator containing 7SK/H1 hybrid promoters, each driving one TDP-43 shRNA (target 1: GAGACTTGGTGGTGCATAA and target 2: GGAGAGGACTTGATCATTA) into the BstBI and SalI sites of pAAV-Puro_siKD^20^. 24 hours post transfection, cells were split into a T150 plate and subjected to selection with 0.75 µg/ml Puromycin (Gibco) for seven days, followed by a 4-day selection with 1.5 µg/ml puromycin. Single colonies were picked and expanded. The inducible TDP-43 knockdown clone was identified by RT-qPCR. Cells were induced with 1 µg/ml doxycycline for 2 days, followed by RNA isolation using the Direct-zol RNA Miniprep Plus kit (Zymo Research) and TDP-43 knockdown was validated by assessment of *TARDBP* mRNA levels by comparing induced and uninduced cells by RT-qPCR with Mesa Green qPCR MasterMix (Eurogentec) according to the manufacturer’s instructions using 40 ng of cDNA and 0.6 µM f.c. primers in a total volume of 20 µl using a RotorGene Q (Qiagen).

### Inducible TDP-43 knockdown SH-SY5Y cells

SH-SY5Y cells containing a doxycycline-inducible short hairpin RNA (shRNA) cassette for TDP-43 are described in ^3^. Cells were maintained in DMEM/F-12 + GlutaMAX (Gibco, 10565018) supplemented with 10% fetal bovine serum (FBS) (Gibco, A5256701) and 1% Penicillin/Streptomycin (Gibco, 15140122) in an incubator maintained at 37°C and 5% CO2. For induction of shRNA against TDP-43, cells were treated with doxycycline hyclate (SigmaAldrich, D9891) at the concentrations and durations detailed for each experiment below.

### Evaluation of U7SmOPT constructs on UNC13A and STMN2 minigenes in TDP-43 depleted 293T cells

To examine the efficiency of the tU7 constructs on cryptic exon splicing for STMN2 and UNC13A, 80% confluent 293T-2xTDP-shRNA cells in 6-well plates were transfected with 200 ng of STMN2 or UNC13A minigenes and 1800 ng U7SmOPT-CMV-BSD plasmids using Mirus TransIT-LT1 (Mirus Bio) according to the manufacturer’s instructions.

For testing the combined targeting of CEs, pMA-3x-U7smOPT containing three tU7 cassettes against STMN2, UNC13A, and INSR was ordered as gene synthesis (GeneArt, LifeTechnologies) and transfections were performed as above, but in the presence of both minigenes. 24 hours post-transfection, cells were split 1:1 and induced with 1ug/ml doxycycline (Sigma Aldrich). 72 hours post-transfection cells were harvested and RNA was isolated using the Absolutely RNA Miniprep Kit (Agilent technologies) according to the manufacturer’s instructions. RNA was reverse transcribed to cDNA using the High-capacity RNA-to-cDNA kit (Applied Biosystems) or LunaScript RT SuperMix Kit (New England BioLabs). TDP-43 mRNA levels as well as the ratio of cryptic to correctly spliced levels of STMN2, UNC13A, and INSR were assessed by RT-qPCR using 20 µl final volume of PowerUp™ SYBR™ Green Master Mix (ThermoFisher) with 40 ng of cDNA, and 0.3 µM primers on a Rotor-Gene Q using the fast cycling mode according to the manufacturer’s instructions. The list of primers used can be found in Table S2 under RT-qPCR primers for minigene testing.

### Lentiviral vector and particle generation

To generate lentiviruses for transduction, U7smOPT strings were cloned into the ClaI sites of a pLVX-EF1a-mCherry T2A-BSD vector. pLVX-EF1a-mCherry T2A-BSD was generated by cloning a gene synthesised string containing the mCherry T2A-BSD ORF between the EcoRI and MluI sites of pLVx-EF1a-IRES-Puro (Clontech Laboratories, Takara Bio) using In-Fusion Snap Assembly EcoDry (Takara Bio) following the manufacturer’s instructions. The U7smOPT cassettes were PCR amplified from their respective U7smOPT-CMV-BSD construct with additional 15 bp overhangs using CloneAmp HiFi PCR (Clonetech Laboratories, Takara Bio) following manufacturer’s instructions with use of 0.3 µM of primers LV inf pLVX Cla forward and LV inf pLVX Cla reverse and 100 ng of template plasmid. This was then cloned into the pLVX-EF1a-mCherryT2A-BSD backbone digested with Cla1 (New England BioLabs) using InFusion Snap Assembly EcoDry (Takara Bio) following the manufacturer’s instructions. 21 µg of the cloned pLVX-EF1a-mCherryT2A-BSD-U7smOPT and 30 µl Trans-Lentiviral Packaging Mix (Dharmacon) was transfected using Lipofectamine 2000 (Invitrogen) following manufacturer’s instructions into >80% confluent HEK293T cells (Takara Bio) cultured in T-150 flasks using DMEM/F12 medium (Gibco) with 10% tetracycline-free FBS and 1% Penicillin/Streptomycin. Medium exchange was performed 24 hours post-transfection and 35ml supernatant was harvested, filtered with 0.45 µm SFCA filter (Thermo Scientific), and supplemented with LentiX Concentrator (1:4, Takara Bio) for the following two days. The mixture was then incubated overnight at 4°C, centrifuged at 1,500 x g for 45 minutes at 4°C, resuspended in a total of 2ml PBS, and aliquoted before flash freezing in liquid nitrogen and storing at –70°C. Virus titre was estimated to be at least 1 x 10^7^ IFU/ml using Lenti-X GoStix (Takara Bio) prior to freezing aliquots.

### Testing of U7smOPT in inducible TDP-43 knockdown SH-SY5Y cells

~2.5 million inducible TDP-43 knockdown SH-SY5Y cells in 5 ml DMEM/F12 medium (Gibco) supplemented with 10% tetracycline-free FBS, 1% Penicillin/Streptomycin and 4 µg/ml Polybrene (SantaCruz) in T-25 flasks were transduced with 100 µl of lentivirus for 24 hours. Stable clones were selected using 2 µg/ml Blasticidin (Gibco) for 4 days followed by 2 days 1 µg/ml Puromycin (Gibco) to reselect stable clones expressing the TDP-43 shRNA cassettes. A T-25 flask was seeded for each line and TDP-43 knockdown was induced the next day by 0.1 µg/ml doxycycline for 5 days and another 5 days of 1 µg/ml doxycycline.

### RT-qPCR and RT-PCR

RNA was extracted from SH-SY5Y cells using the Absolutely RNA Miniprep Kit (Agilent technologies) according to the manufacturer’s protocol and first strand cDNA synthesis was performed with LunaScript RT SuperMix Kit (New England BioLabs).

PCR was then conducted by amplifying strands using Platinum SuperFi II DNA Polymerase (ThermoFischer) and resolved on 3% agarose gel with 1:10’000 PAGE GelRed® Nucleic Acid Gel Stain (Biotium). Correct and cryptic *UNC13A* and *STMN2 transcripts*, as well as *TARDBP* mature mRNA levels were quantified by RT-qPCR using 20 µl final volume of PowerUp™ SYBR™ Green Master Mix (ThermoFisher) with 40 ng of cDNA and 0.3 µM primers on a RotorGene Q using the fast-cycling mode according to the manufacturer’s instruction. Data was analysed using the ΔΔCt method^21^ and normalised to *GAPDH*. Primers used are listed in **Table S2**.

### iPSC cell culture

The WTC11 human iPSCs^22^ used in this study were previously engineered to overexpress human neurogenin-2 (*NGN2*) to generate cortical-like i3Neurons, or an hNIL construct expressing human transcription factors Islet-1 (*ISL1*) and LIM Homeobox 3 (*LHX3*), and *NGN2*, to generate i3-lower motor neurons (i3-LMNs), under a doxycycline-inducible promoter, alongside an expression cassette for an enzymatically dead Cas9 (+/− CAG-dCas9BFP-KRAB), are previously described^23-25^. The generation of the Halo-TDP line is described in^6^. iPSCs were maintained in Essential 8 Flex medium (Gibco, A2858501) with daily feeds and kept in an incubator at 37°C and 5% CO2. Cells were routinely passaged twice-weekly using Versene (Gibco, 15040066) in Geltrex-coated (1:100; Gibco, A14133-01) tissue culture dishes. Chromosome analysis was performed periodically to exclude a karyotypic abnormality, and cell pellets were tested for mycoplasma contamination.

### Generation of iPSC lines stably expressing tU7s

tU7 iPSC lines were generated analogously to tU7 SH-SY5Y by lentiviral transduction with the following modifications. A cell suspension of 250,000 iPSCs, generated by Accutase (Gibco, A1110501) in Essential 8 Flex medium (Gibco, A2858501) treatment, was transduced with 50ul lentiviral vectors overnight in the presence of 10 mg/ml Polybrene (Sigma, H9268) in one well of a 12-well plate. The following morning, cells were washed with PBS and culture medium was replaced. Two days after lentiviral delivery, cells were selected for 72 hours with 10 mg/ml Blasticidin (Sigma, SBR000221ML), followed by expansion of cell pools.

### Differentiation of iPSCs into cortical-like neurons (i3Neurons)

The differentiation process was as reported previously^6,23^. Briefly, to initiate neuronal differentiation, 2.5 million iPSCs per 10 cm plate were dissociated using Accutase (Gibco, A1110501) on day 0 and re-plated onto Geltrex-coated (1:100; Gibco, A14133-01) tissue culture dishes in induction medium: DMEM/F-12 + GlutaMAX medium (Gibco, 31331028), 1x MEM Non-Essential Amino Acids (Gibco, 1140050), 2 μg/ml Doxycycline hyclate (SigmaAldrich, D9891), 1x N2 supplement (Gibco, 17502048), 2 μM XAV939 (Cambridge Bioscience, SM38-10), 10 μM SB431542 (Biotechne, 1614), and 100 nM LDN-193189 (Cambridge Bioscience, 19396-5mg-CAY). 1x ROCK Inhibitor Y-27632 dihydrochloride (Tocris, 1254/10) was added to the medium on the day of plating. Culture medium was changed daily during this stage.

On day 3, pre-neuron cells were re-plated onto dishes coated with 50 μg/ml Poly-D-Lysine (Gibco, A38904-01) and 10 μg/ml Laminin (Gibco, 23017015) in either 96-well plates (12,50025,000 cells per well) for Incucyte® Live-Cell Analysis System experiments, or 12-well dishes (500,000 cells per well) for RNA and protein extraction. They were plated in i3Neuron cortical neuron culture medium: BrainPhys neuronal culture medium (StemCell Technologies, 5790), supplemented with 1x B27 (Gibco, 17504044), 1x N2 (Gibco, 17502048), 10 ng/ml BDNF (PeproTech, 450-02), 10 ng/ml GDNF (PeproTech, 450-10), and 1 μg/ml Laminin (Gibco, 23017015). 1x RevitaCell (containing ROCK inhibitor to facilitate single-cell survival; Gibco, A2644501) was added on the day of plating. 24 hours after plating, culture medium was fully replaced to remove RevitaCell, except for Incucyte® experiments where it was kept in owing to ROCK inhibition being known to promote neurite outgrowth. Following this, i3Neurons were fed twice weekly by half-medium changes. For PROTAC-mediated knockdown of Halo-TDP43, i3Neurons were treated with 300 nM HaloPROTAC. HaloPROTAC was added from the first day of differentiation (day 0) for Incucyte® neurite outgrowth experiments, and from day 4 for RNA and protein analysis experiments.

For electrophysiology experiments, i3Neurons were maintained in neuron media: BrainPhys supplemented with 1x N2 supplement, 1x B27 supplement, 10 ng/ml NT3 (Stemcell Technologies, 78074.1), 20 ng/mL BDNF, 20 ng/ml GDNF, 2 mM dibutyrl cAMP (Sigma, D0627), 200 nM L-ascorbic acid (Sigma, A0278), and 1 μg/mL laminin and half media changes were performed twice per week.

### Differentiation of iPSCs into lower motor neurons (i3-LMNs) and CRIPSRi

For induction of WTC hNIL dCas9BFP-KRAB iPSCs to i3LMN, iPSCs were plated in induction medium containing DMEM/F-12 + GlutaMAX medium (Gibco, 31331028), MEM Non-Essential Amino Acids (Gibco, 1140050), N2 supplement (Gibco, 17502048), 0.2 mM compound E, 2 μg/ml Doxycycline hyclate (Sigma-Aldrich, D9891), 1 μg/ml Laminin (Thermo, 23017015). After induction for 48 hours, differentiated motor neurons were re-plated onto dishes coated with 50 μg/ml Poly-D-Lysine (Gibco, A38904) and 10 μg/ml Laminin in induction medium supplemented with 1 μg/ml Laminin and 10 μM ROCK inhibitor Y-27632 dihydrochloride (Tocris, 1254/10). After 24 hours, induction medium was replaced with motor neuron medium containing Neurobasal (Gibco, 21103049), N2 max supplement (R&D Systems, AR009), N21 max supplement (R&D Systems, AR008), CultureOne supplement (antimitotic agent; Gibco, A3320201), 1 μg/ml Laminin, 2 μg/ml Doxycycline hyclate, 10 ng/mL BDNF (Peprotech) and 10 ng/ml GDNF (Peprotech). Half-medium changes were performed twice a week. Cells were maintained in an incubator at 37°C and 5% CO2. CRISPRi-mediated depletion of TDP-43 was performed as described in^3,6^.

### RNA extraction, cDNA synthesis, RT-PCR and qPCR from iPSC-derived neurons

RNA extraction from cells was carried out using the RNeasy® mini kit (Qiagen, 74104) or the Direct-zol RNA MiniPrep kit (Zymo Research, R2051) following the manufacturer’s protocol. RNA was eluted in 30 μl RNase-free water and concentrations were quantified using a Nanodrop. RNA was stored at −80°C until use.

250-1,000 ng of RNA was used per reaction for reverse transcription (RT). Samples for RNAsequencing were assessed for RNA quality on a TapeStation 4200 (Agilent), and bands were quantified with TapeStation Systems Software v3.2 (Agilent). First-strand cDNA synthesis was performed using the RevertAid kit (Thermo Scientific, K1622) following the manufacturer’s protocol. cDNA was diluted to 5-10 ng/μl and stored at −20°C until use. For PCR, cDNA was amplified using the Q5® High-Fidelity DNA polymerase (New England Biolabs, M0491). PCR products were resolved on a TapeStation 4200 (Agilent).

Gene expression analysis was performed by PCR using Taqman Multiplex Master Mix (Thermo, 4461882), or Power Up SYBR Green Master Mix (Thermo, A25742). Transcript levels were quantified on a QuantStudio 5 Real-Time PCR system (Applied Biosystems) using the ΔΔCt method^21^. Using RefFinder, we identified GAPDH as the most stable endogenous control across our conditions of interest. Primers and assays for RT-PCR and RT-qPCR are described in **Table S2**.

### Western blotting (iPSC-derived neurons)

Cells were lysed either directly in NuPage 4x LDS Sample Buffer (Invitrogen, NP0007) or in RIPA buffer (Thermo Scientific, 89900) and protein concentration was measured using the Pierce 660 nm Protein Assay Kit (Thermo Scientific, 22662). Lysates were heated at 95 °C for 5 min with 100 mM dithiothreitol (DTT reducing agent; Thermo Scientific, R0861) or 1x NuPage LDS Sample buffer and 2.5% beta-mercaptoethanol to denature the protein, reduce disulphide bonds, and ensure proper separation and detection. Lysates were passed through a QIAshredder (Qiagen, 79656) to homogenise the protein. Equal amounts of protein lysates were loaded alongside a ladder of appropriate size and resolved on 4-12% Bis-Tris Gels (Invitrogen, NP0321BOX) with MOPS buffer (Invitrogen, NP0001), at a constant voltage of 120 V for 90-180 minutes, depending on the separation of bands required. Proteins were then transferred to 0.2 μm PVDF membranes (Amersham GE Healthcare, GE10600057) in NuPAGE Transfer Buffer (Invitrogen, NP0006) at a constant current of 0.2 A for 90-180 minutes, depending on protein size or in Trans-Blot Turbo 5x Transfer Buffer (BioRad, 10026938) using the Trans-Blot Turbo Transfer System for 30 minutes. After blocking with 5% milk for 1 hour, blots were probed with primary antibodies (rabbit anti-STMN2, ProteinTech 10586-1-AP, 1:1000; rabbit anti-UNC13A, Synaptic Systems 126103, 1:2000; mouse antiTDP-43, abcam ab104223 clone 3H8, 1:5000; rat anti-tubulin, Millipore MAB1864 clone YL1/2, 1:5000) at 4°C overnight. After three 10-minute TBS-T (Thermo Scientific, J77500) washes, blots were probed with HRP-conjugated secondary antibodies (goat anti-rabbit, Bio-Rad 1706515, 1:10000; goat anti-mouse, BioRad 1706516, 1:10000; goat anti-rat, Dako P0450, 1:10000 or Ab6845, 1:10000) for 1 hour at room temperature. After three further 10-minute TBS-T washes, western blots were developed with Chemiluminescent HRP substrate (Merck Millipore, WBKLS0500) on a Bio-Rad ChemiDoc Imaging System and quantified using ImageJ software. All protein quantifications were normalised to Tubulin levels.

### Neurite outgrowth assay

After three days of induction medium, Halo-TDP-43 i3Neurons stably expressing either a nontargeting tU7 Ctrl construct, or a *STMN2*-targeting bifunctional tU7 construct, were plated in i3Neuron cortical culture medium with RevitaCell supplement (Gibco, A2644501) in a 96-well plate coated with Poly-D-lysine (Gibco, A3890401) and Laminin (Gibco, 23017015). There were four conditions: i) tU7 Ctrl-expressing i3Neurons without HaloPROTAC treatment; tU7 Ctrl-expressing i3Neurons with HaloPROTAC treatment; tU7 STMN2-expressing i3Neurons without HaloPROTAC treatment; and tU7 STMN2-expressing i3Neurons with HaloPROTAC treatment. For each condition, two cell densities were plated (8,000 and 12,000 cells). For each density, eight wells were plated, comprising eight technical replicates per condition per density. In conditions where TDP-43 knockdown was desired, HaloPROTAC treatment was added to the medium (300 nM; Promega, GA3110) from the first day of induction medium. One hour after plating i3Neurons in a 96-well plate, the plate was placed in an Incucyte® (Sartorius) Live-Cell Analysis System for longitudinal visualisation and analysis of neurite outgrowth. The Incucyte® machine is a real-time live-cell analysis system that allows monitoring and analysis of cell behaviour over time, capturing high-resolution images of cells in culture within a standard tissue culture incubator machine maintained at 37°C and 5% CO2. The machine was set up to capture four images within each well, every two hours, for a duration of 48 hours. Brightfield, as well as red fluorescence images (given the mCherry reporter in the tU7 constructs) were captured. Of note, RevitaCell supplement containing ROCK inhibitor was kept in the medium from plating and throughout the imaging period, given its known effect on neurite outgrowth, to avoid introducing fluctuations caused by removing it. Five independent differentiations were performed, comprising five biological replicates. Using cell body and neurite masks on the Incucyte® software, cell body area (μm^2^) and neurite length (mm/mm2) were quantified in an automated way, ensuring that the masks had a good fit, and a mean and SEM were calculated for each condition and each timepoint from all technical replicates. One of the two plating densities was selected for each condition, based on the average cell body area for identification of which densities were most similar between conditions. To account for any variation in cell densities between conditions, mean neurite length was normalised to mean cell body area by dividing the former by the later. The neurite length/cell body area value was normalised to the first time point by dividing the value at each timepoint by the value for the first time point, such that each condition started at a value of 1.00 at the first time point. This normalised value was plotted over time for each of the four conditions. Data from each differentiation were first plotted on separate graphs. Further, for each biological replicate, a delta value was then calculated for each U7-expressing iPSC line by subtracting the values for neurite length/cell body area normalised to the first timepoint with HaloPROTAC treatment, from the values without HaloPROTAC treatment. This was performed for each timepoint, and a mean and SEM of the delta value from all replicates was plotted for each U7-expressing cell line.

### UNC13A and Synapsin immunofluorescence

500,000 cells were grown on 50,000 primary rat astrocytes on Poly-D-Lysine/Laminin-coated coverslips. Neurons were fixed at day 35 in 4% PFA for 10 minutes. Cells were then permeabilised in 0.1% Triton X-100 in PBS for 10 minutes and washed three times in PBS. Cells were blocked in 2% BSA in PBS for one hour. Primary antibodies, guinea pig antiMunc13–1 (Synaptic Systems, 126–104) and mouse anti Synapsin 1 (Synaptic Systems, 106– 011), were diluted in PBS at 1:500 and 1:1,000, respectively, and cells were incubated in primary antibody for one hour at room temperature. Cells were washed three times in PBS. Secondary antibodies, goat anti-mouse IgG (H+L) AlexaFluor 488 (Thermo) and goat antiguinea-pig IgG (H+L) AlexaFluor 647 (Thermo), were diluted in PBS 1:1,000, and cells were incubated in secondary antibody at room temperature for one hour. Cells were further washed three times in PBS and mounted in Mowiol-4–88 (PolySciences, 17951-500) mounting medium containing 1:1,000 DAPI (Invitrogen, D1306). Cells were imaged using a Zeiss 980 Airyscan confocal at 63x with a 3x crop. Z-stacks of 7× 0.5 μm intervals were taken. Fiji was used to analyse mean UNC13A intensity in an automated way based on manually selected synapsin-positive ROIs.

### Electrophysiology experiments

For whole-cell patch clamp recordings, experiments were performed as previously described by our lab^6^. 500,000 cells were plated onto 18 mm No.1 coverslips (Deckglaser) coated with 50 μg/ml Poly-D-Lysine (Gibco, A3890401) and 10 μg/l Laminin (Gibco, 23017015). Primary astrocytes, obtained from E18 rat cortices via 15-minute trypsinisation (10 mg/ml; Gibco), were plated one week prior and treated with CultureOne supplement (Gibco, A3320201) two days prior. Miniature excitatory post-synaptic currents (mEPSCs) were recorded at four-weeks maturation on rat astrocytes, in extracellular solution containing: 136 mM NaCl, 2.5 mM KCl, 10 mM HEPES, 1.3 mM MgCl2, 2 mM CaCl2, 10 mM Glucose, pH 7.3, 300 mOsm, supplemented with 10 μM tetrodotoxin and 10 μM gabazine. Patching pipettes were filled with cesium methanosulfate based internal solution: 135 mM CsMeSO3, 10 mM HEPES, 10 mM Na2-Phosphocreatine, 5 mM Glutathione, 4 mM MgCl2, 4 mM Na2ATP, 0.4 mM NaGTP supplemented with 5 μM QX-314. Pipettes were pulled from borosilicate glass (O.D. 1.5 mm, I.D. 0.86 mm, Sutter instruments) to a resistance of between 3-5 MΩ. Whole-cell patch clamp recordings were then made without whole-cell compensation applied for 5 minutes at the soma. Cells were held at −70 mV using a Multiclamp 700B amplifier (Molecular Devices) and data acquired using a Digidata 1440A digitiser (Molecular Devices). All recordings were carried out on a heated stage set to 37°C. Data were acquired with Clampex software (Molecular Devices) and Axon Multiclamp Commander Software (Molecular Devices), with data sampled at a rate of 20 kHz and filtered at 10 kHz. mEPSCs were analysed in a fully automated manner using Clampfit 10.7 software (Molecular devices).

### Generation of pAAV vectors and virus production

To generate pAAV expressing shRNA and U7 snRNA constructs, the H1 promoter, hairpin and shRNA sequences (shCtrl: 5’-ACGTGACACGTTCGGAGATAA-3’ and shTardbp: 5’ TTTCCAACAATGACAACTCTT-3’) were cloned into pAAV EGFP U7smOPT bifunctional constructs using Acc65I restriction sites. rAAVPHP.eB viruses were produced as previously described^26^. AAV vectors were co-transfected with helper plasmids in HEK293T cells using polyethylenimine (Polysciences, Inc.) and then harvested 72 hours following transfection. The cells were lysed in the presence of 0.5% sodium deoxycholate and 50 Units/mL Benzonase (Sigma-Aldrich) by several cycles of freeze-thaw and viruses were then isolated using a discontinuous iodixanol gradient. The viral titter was measured by qRT-PCR and virus was then diluted in sterile phosphate-buffered saline (PBS).

### Animal studies

All procedures relating to the use of animals were conducted in accordance with the National Institutes of Health Guide for Care and Use of Experimental Animals and were approved by the Mayo Clinic Institutional Animal Care and Use Committee (IACUC Protocol number A00007254-23).

### Neonatal viral injections

Intracerebroventricular (ICV) injections of virus were performed as previously described^26^. Briefly, 2 μl (1.5E10genomes/μl) of AAV PHP.eB-shCtrl;U7 Stmn2, -shTardbp;U7 Ctrl or shTardbp;U7 Stmn2, were diluted in sterile saline and injected into the lateral ventricles of cryoanesthetized humanized Stmn2+/+ (hStmn2+/+, C57BL/6J-Stmn2em6(STMN2)Lutzy/Mmjax, MMRRC stock #069791-JAX; Jackson Laboratories)^10^ mouse pups on postnatal day 0 (P0). Pups were allowed to recover following the injection on a heating pad and then returned to the home cage with their mother. Confirmation of mouse genotype was performed as described by Jackson Laboratories using a common forward primer and reverse primers specific to the wild-type or hStmn2 allele and are listed in **Table S2**.

### Tissue processing

Mice were euthanized by carbon dioxide and brains and spinal cords were harvested. The brain was cut across the midline, and the left half was fixed in 4% paraformaldehyde, while the other half of the brain was dissected into forebrain and cerebellum and flash frozen. In total brain and spinal cord from 21 shCtrl; U7 Stmn2 (7 male and 14 female), 9 shTardbp; U7 Ctrl (2 male and 7 female) and 11 shTardbp; U7 Stmn2 (5 male and 6 female) mice were collected.

### RNA extraction, reverse transcription and qPCR from tissue samples

For RNA extraction, forebrains were first homogenized in Tris EDTA buffer, pH 8.0, containing protease and phosphatase inhibitors. Homogenates were mixed with TRIzol LS (Thermo Fisher), and total RNA was extracted using the Direct-zol RNA MiniPrep kit (Zymo Research) according to manufacturer’s instructions. cDNA was obtained following reverse transcription of extracted RNA using the High-Capacity Transcription Kit (Applied Biosciences). Analysis of RNA was performed using qRT-PCR. Each sample was measured in triplicate with the SYBR green assay (Life Technologies) on the ABI Prism 7900HT Fast Real-Time PCR System (Applied Biosystems). Data was analyzed using the ΔΔCt method and normalized to endogenous controls Gapdh and Rplp0. The sequences of the primers used for this study are listed in **Table S2**.

### Preparation of tissue lysates and western blotting

Homogenates were lysed in co-immunoprecipitation buffer (50 mM Tris–HCl, pH7.4, 300 mM NaCl, 1% Triton X-100, 5 mM EDTA, 2% SDS) with protease and phosphatase inhibitors, sonicated on ice, and centrifuged at 16,000 x g for 15 minutes. Supernatants were moved to a new tube and protein concentration of lysates was determined by BCA assay (Thermo Fisher Scientific). Samples were prepared by mixing cell lysates with Tris-Glycine SDS sample buffer (ThermoFischer Scientific) and 5% beta-mercaptoethanol (Sigma Aldrich) and then heat denatured at 95°C for 5 minutes. Samples were run on 4–20% Tris-glycine gels and transferred onto PVDF membranes (Millipore). Following transfer, membranes were blocked with 5% non-fat dry milk in TBS plus (Tris Buffered Saline with 0.1% Triton X-100, TBS-T) for 1 hour, and then incubated with primary antibody overnight at 4°C to probe for Tdp-43 (1:5000, 12892-1-AP, Proteintech), Stmn2 (1:2000, 10586-1-AP, Proteintech), or Gapdh (1:10000, H86504M, Meridian). Membranes were washed in TBS-T and incubated with appropriate secondary antibodies conjugated to horseradish peroxidase (1:5000; Jackson ImmunoResearch) for 1 hour. After washing in TBS-T, protein expression was visualized by enhanced chemiluminescence (PerkinElmer) and exposure to film. Quantitative densitometry was performed using ImageJ, and Tdp-43 and Stmn2 levels were normalized to Gapdh expression, and to then to the levels of control group (shCtrl; U7 Stmn2).

