## Supplementary Information for "U7 small nuclear RNA splice-switching therapeutics for STMN2 and UNC13A in Amyotrophic Lateral Sclerosis"

#### Supplementary Information Guide

| | |
|---|---|
| **Supplementary figure 1:** | tU7 screening information |
| **Supplementary figure 2:** | Additional quantification of transcript and protein changes following tU7 Treatment in neurons and humanized Mice |
| **Supplementary table 1:** | U7 SmOPT snRNA sequences |
| **Supplementary table 2:** | Primer sequences/qPCR assay information |

#### Supplementary Figure S1

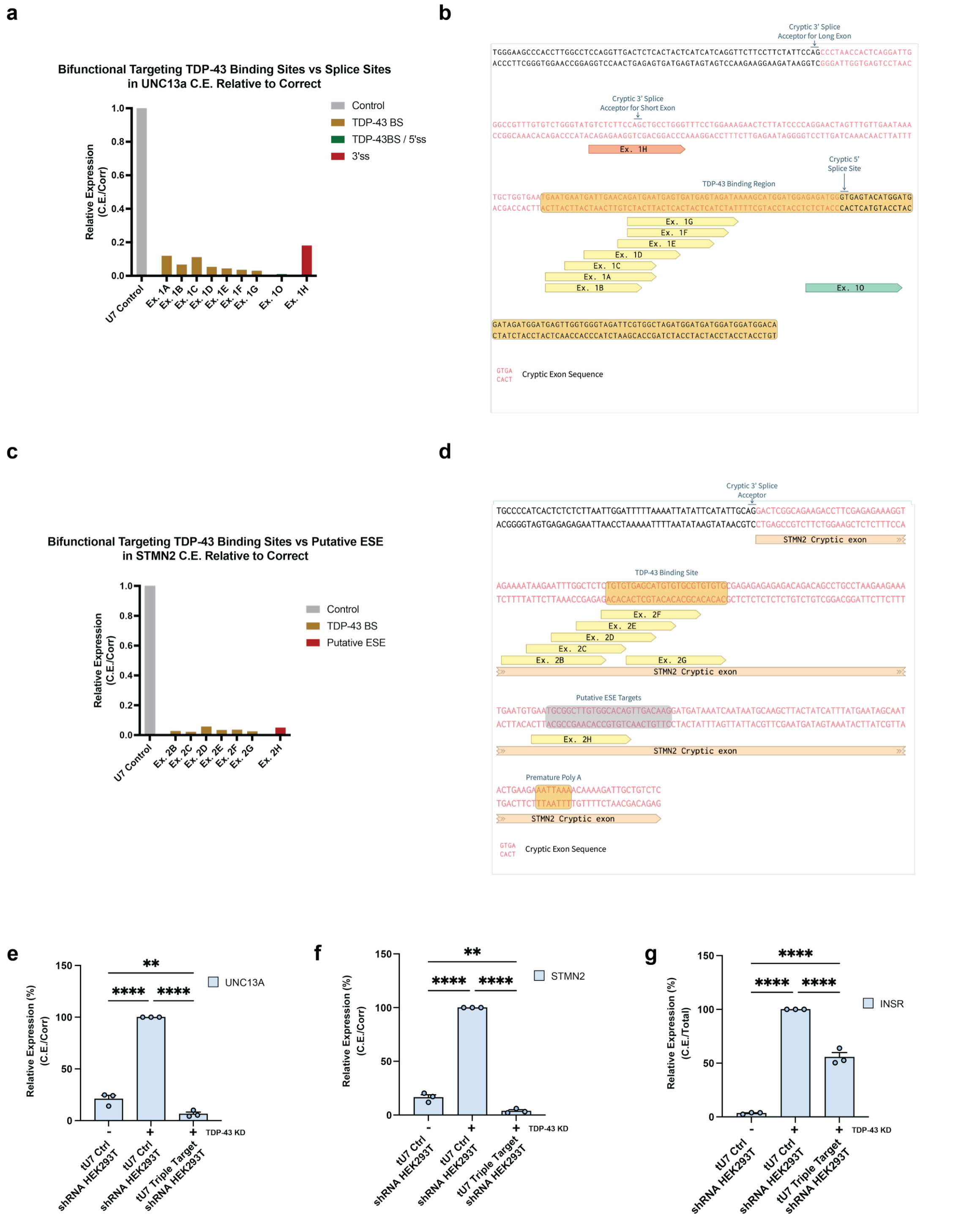


**Supplementary Figure 1: a, c** Ratio of cryptic exon splicing to correct splicing of the UNC13A (**a**) and STMN2 (**c**) minigene in the presence of different modified U7 snRNA constructs containing either an antisense sequence that targets the 5'-splice site, TDP-43 binding region, 3'-splice site or putative exonic splicing enhancer (ESE) tested in TDP-depleted 293T cells with UNC13A or STMN2 minigene. *n* = 1. **b, d** Shows the TDP-43 regulated UNC13A (**b**) and STMN2 (**d**) cryptic exon target and flanking regions annotated with splicing elements and tU7 antisense binding regions for screened constructs. **e, f, g** Ratio of cryptic to correctly or total (mean ± SEM)

transcript RT-qPCR levels of *UNC13A* (**e**), *STMN2* (**f**) and *INSR* (**g**) mature mRNA in the TDP-depleted 293T cells transfected with STMN2 and UNC13A minigenes upon transfection with non-targeting control (tU7 Ctrl) or a combined vector comprising multiple U7 constructs pMA-3x-U7SmOPT (tU7 Triple Target). The 3x-tU7SmOPT construct contains three U7 constructs (Ex.1O, Ex. 2C, tU7 INSR, see Table S1) in tandem each comprising a different antisense sequence complementary to UNC13A, STMN2 and INSR cryptic exons. Results are normalised to tU7 Ctrl without TDP-43 KD. n=3. Statistical significance was determined by one-way ANOVAs with Tukey multiple comparison (*$p < 0.05$; **$p < 0.01$; ***$p < 0.001$; ****$p < 0.0001$).

#### Supplementary Figure S2

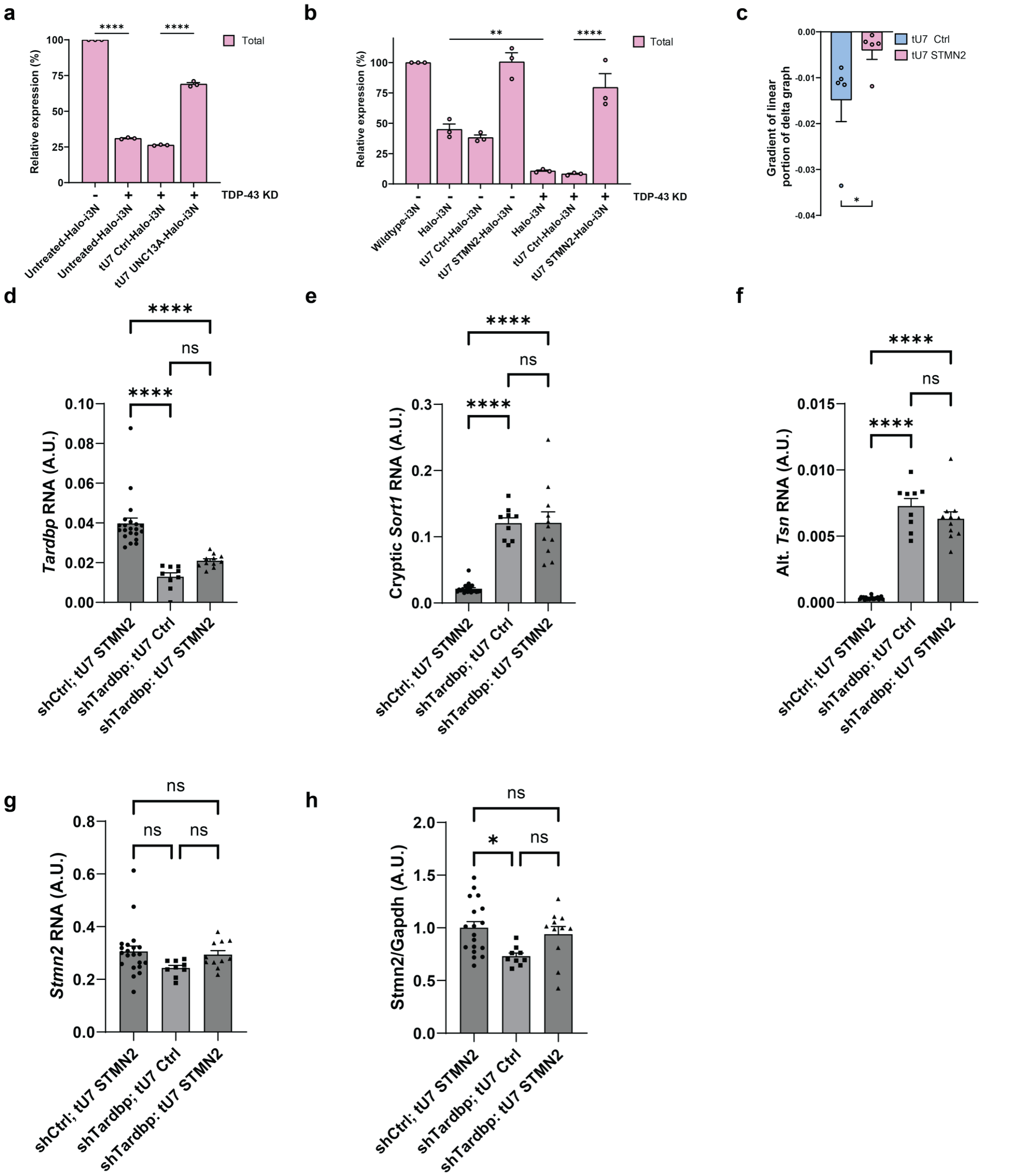


**Supplementary Figure 2: a,** Quantification (mean ± SEM) of RT-qPCR levels of total *UNC13A* **(a)** and *STMN2* **(b)** transcripts shows successful rescue after respective tU7 treatment. *n* = 3. **c,** Quantification (mean ± SEM) of the gradient of the delta curves' linear portion in **Fig. 2n** for tU7 Ctrl and tU7 STMN2-expressing i3Neurons, showing that the tU7 STMN2, rescued TDP-43 loss-dependent neurite outgrowth impairment. *n* = 5. **d-h,** RNA measured form the forebrains of 1.5-month-old humanized Stmn2$^{+/+}$ mice ICV-injected with AAV expressing shRNA and tU7 snRNA, *shCtrl;* tU7 STMN2 (*n* = 18-21), *shTardbp;* tU7 Ctrl (*n* = 9), and *shTardbp;* tU7 STMN2 (*n* = 11). d, *Tardbp* RNA, normalised to endogenous controls *Rplp0* and *Gapdh*, as measured by RT-qPCR, was significantly reduced in *shTardbp* compared to s*hCtrl*-treated mice. **e-f,** Cryptic *Sort1* **(e)**

and alternatively spliced Tsn RNA **(f)**, normalised to endogenous controls *Rplp0* and *Gapdh*, as measured byRT-qPCR, were significantly increased in *shTardbp* compared to s*hCtrl*-treated mice. **g,** *Stmn2* RNA, normalised to endogenous controls *Rplp0* and *Gapdh*, as measured by RT-qPCR, was not significantly different in any groups. **h,** Stmn2 protein normalised to Gapdh levels are significantly reduced in *shTardbp*; tU7 Ctrl conditions, but similar in *shTardbp*; tU7 STMN2 compared to *shCtrl*; tU7 STMN2-treated mice. Statistical significance for **a,b, and d-h** was determined by one-way ANOVA followed by Tukey's multiple comparisons test, and for **c** was determined by performing a one-tailed two-sample Wilcoxon signed rank test (*$p < 0.05$; **$p < 0.01$; ***$p < 0.001$; ****$p < 0.0001$; ns = non-significant).

### Table S1: U7 SmOPT snRNA sequences

| Name | Target Antisense Sequence (Sequences taken forward in bold) | tU7 smOPT snRNA Sequence (hnRNPA1 tail underlined) |
|---|---|---|
| tU7 Ctrl | GTGTTACAGCTCTTTTAG | AAUAUGAUAGGGACUUAGGGUGGUGUUACAGCUCUUUUAGAAUUUUUGGAGCAGGUUUUCUGACUUCGGUCGGAAAACCCCU |
| Ex. 1A | TTCATCTGTTCAATCATTCATTC | AAUAUGAUAGGGACUUAGGGUGUUCAUCUGUUCAAUCAUUCAUUCAAUUUUUGGAGCAGGUUUUCUGACUUCGGUCGGAAAACCCCU |
| Ex. 1B | ATCTGTTCAATCATTCATTC | AAUAUGAUAGGGACUUAGGGUGAUCUGUUCAAUCAUUCAUUCAAUUUUUGGAGCAGGUUUUCUGACUUCGGUCGGAAAACCCCU |
| Ex. 1C | TTCATCTGTTCAATCATTC | AAUAUGAUAGGGACUUAGGGUGUUCAUCUGUUCAAUCAUUCAAAUUUUUGGAGCAGGUUUUCUGACUUCGGUCGGAAAACCCCU |
| Ex. 1D | ACTCATTCATCTGTTCAATC | AAUAUGAUAGGGACUUAGGGUGACUCAUUCAUCUGUUCAAUCAAUUUUUGGAGCAGGUUUUCUGACUUCGGUCGGAAAACCCCU |
| Ex. 1E | ACTCATCACTCATTCATCTG | AAUAUGAUAGGGACUUAGGGUGACUCAUCACUCAUUCAUCUGAAUUUUUGGAGCAGGUUUUCUGACUUCGGUCGGAAAACCCCU |
| Ex. 1F | TCTACTCATCACTCATTCATC | AAUAUGAUAGGGACUUAGGGUGUCUACUCAUCACUCAUUCAUCAAUUUUUGGAGCAGGUUUUCUGACUUCGGUCGGAAAACCCCU |
| Ex. 1G | TATCTACTCATCACTCATTCATC | AAUAUGAUAGGGACUUAGGGUGUAUCUACUCAUCACUCAUUCAUCAAUUUUUGGAGCAGGUUUUCUGACUUCGGUCGGAAAACCCCU |
| **Ex. 1O** | **TCCATGTACTCACCCATCTC** | **AAUAUGAUAGGGACUUAGGGUGUCCAUGUACUCACCCAUCUCAAUUUUUGGAGCAGGUUUUCUGACUUCGGUCGGAAAACCCCU** |
| Ex. 1H | CCCAGGCAGCTGGAAGAGAC | AAUAUGAUAGGGACUUAGGGUGCCCAGGCAGCUGGAAGAGACAAUUUUUGGAGCAGGUUUUCUGACUUCGGUCGGAAAACCCCU |
| Ex. 2B | GAGAGCCAAATTCTTATTTTC | AAUAUGAUAGGGACUUAGGGUGGAGAGCCAAAUUCUUAUUUUCAAUUUUUGGAGCAGGUUUUCUGACUUCGGUCGGAAAACCCCU |
| **Ex. 2C** | **CACAGAGAGCCAAATTCTTA** | **AAUAUGAUAGGGACUUAGGGUGCACAGAGAGCCAAAUUCUUAAAUUUUUGGAGCAGGUUUUCUGACUUCGGUCGGAAAACCCCU** |
| Ex. 2D | TGCTCACACAGAGAGCCAAAT | AAUAUGAUAGGGACUUAGGGUGUGCUCACACAGAGAGCCAAAUAAUUUUUGGAGCAGGUUUUCUGACUUCGGUCGGAAAACCCCU |
| Ex. 2E | CACATGCTCACACAGAGAGC | AAUAUGAUAGGGACUUAGGGUGCACAUGCUCACACAGAGAGCAAUUUUUGGAGCAGGUUUUCUGACUUCGGUCGGAAAACCCCU |
| Ex. 2F | ACGCACACATGCTCACACAG | AAUAUGAUAGGGACUUAGGGUGACGCACACAUGCUCACACAGAAUUUUUGGAGCAGGUUUUCUGACUUCGGUCGGAAAACCCCU |
| Ex. 2G | CACACACGCACACATGCTCA | AAUAUGAUAGGGACUUAGGGUGCACACACGCACACAUGCUCAAUUUUUGGAGCAGGUUUUCUGACUUCGGUCGGAAAACCCCU |
| Ex. 2H | CTGTGCCACAAGCCGCATTC | AAUAUGAUAGGGACUUAGGGUGCUGUGCCACAAGCCGCAUUCAAUUUUUGGAGCAGGUUUUCUGACUUCGGUCGGAAAACCCCU |
| Ex. 2I | CGAAGGTCTTCTGCCGAGTCCTGC | AAUAUGAUAGGGACUUAGGGUGCGAAGGUCUUCUGCCGAGUCCUGCAAUUUUUGGAGCAGGUUUUCUGACUUCGGUCGGAAAACCCCU |
| **tU7 INSR** | **CCCGTATCCGGTACTATATG** | **AAUAUGAUAGGGACUUAGGGUGCCCGUAUCCGGUACUAUAUGAAUUUUUGGAGCAGGUUUUCUGACUUCGGUCGGAAAACCCCU** |

### Table S2: Primer sequence/qPCR assay information

| **Primers for validation of HEK 293T 2x shTDP cell line** | | |
|---|---|---|
| **Target / Section** | **Primer** | **Sequence (5'-3')** |
| Sybr *TARDBP* | Forward | AACCGAACAGGACCTGAAAGAG |
| Sybr *TARDBP* | Reverse | CAGTCACACCATCGTCCATCTATC |
| Sybr *ACTB* | Forward | TCCATCATGAAGTGTGACGT |
| Sybr *ACTB* | Reverse | TACTCCTGCTTGCTGATCCAC |
| **Primers used in HEK 293T 2x shTDP minigene assays** | | |
| **Target / Section** | **Primer** | **Sequence (5'-3')** |
| *TARDBP* | Forward | TCATCCCCAAGCCATTCAGG |
| *TARDBP* | Reverse | TGCTTAGGTTCGGCATTGGA |
| *GAPDH* | Forward | CCAGAACATCATCCCTGCCT |
| *GAPDH* | Reverse | GGTCAGGTCCACCACTGACA |
| Canonical *UNC13A* | Forward | ACCTGTCTGCATGAGAACCT |
| Canonical *UNC13A* | Reverse | GGGCTGTCTCATCGTAGTAAAC |
| Cryptic *UNC13A* | Forward | ATGGATGGAGAGATGGAACCT |
| Cryptic *UNC13A* | Reverse | GGGCTGTCTCATCGTAGTAAAC |
| Canonical *STMN2* | Forward | GCTAAAACAGCAATGGCCTAC |
| Canonical *STMN2* | Reverse | TTGCTTCACTTCCATATCATCG |
| Cryptic *STMN2* | Forward | GCTAAAACAGCAATGGGACTC |
| Cryptic *STMN2* | Reverse | GCAGGCTGTCTGTCTCTCTC |
| Total *INSR* | Forward | TGGGACCGCTTTACGCTTC |
| Total *INSR* | Reverse | GAGACTGGCTGACTCGTTGAC |
| Cryptic *INSR* | Forward | CTCTGGGACTGGAGCAAAC |
| Cryptic *INSR* | Reverse | CATCCCGTATCCGGTAAGG |
| **Primers used in SH-SY5Y models** | | |
| **Target / Section** | **Primer** | **Sequence (5'-3')** |
| *UNC13A* RT-PCR | Forward | CCGTACCATGTCCAGTACAC |
| *UNC13A* RT-PCR | Reverse | AGTAAACCTTCCAGGCATCG |
| *STMN2* RT-PCR | Forward | GCTCTCTCCGCTGCTGTAG |
| *STMN2* RT-PCR | Reverse | CGAGGTTCCGGGTAAAAGCA |
| *STMN2* RT-PCR | Cryptic reverse | CTGTCTCTCTCTCTCGCACA |
| *TARDBP* | Forward | TCATCCCCAAGCCATTCAGG |
| *TARDBP* | Reverse | TGCTTAGGTTCGGCATTGGA |
| *GAPDH* | Forward | CCAGAACATCATCCCTGCCT |
| *GAPDH* | Reverse | GGTCAGGTCCACCACTGACA |
| Correct *UNC13A* | Forward | ACCTGTCTGCATGAGAACCT |
| Correct *UNC13A* | Reverse | GGGCTGTCTCATCGTAGTAAAC |
| Cryptic *UNC13A* | Forward | ATGGATGGAGAGATGGAACCT |
| Cryptic *UNC13A* | Reverse | GGGCTGTCTCATCGTAGTAAAC |
| Canonical *STMN2* | Forward | GCTAAAACAGCAATGGCCTAC |
| Canonical *STMN2* | Reverse | TTGCTTCACTTCCATATCATCG |
| Cryptic *STMN2* | Forward | GCTAAAACAGCAATGGGACTC |
| Cryptic *STMN2* | Reverse | GCAGGCTGTCTGTCTCTCTC |
| **Primers for cloning pLVX-EF1a-mCherryT2A-BSD-U7smOPT plasmid** | | |
| **Target / Section** | **Primer** | **Sequence (5'-3')** |
| LV inf pLVX Cla | Forward | AGATCCAGTTTATCGATACCAACATAGGAGCTGTGATTGG |
| LV inf pLVX Cla | Reverse | ATGAATTACTCATCGGCGAGAAAGGAAGGGAAGAAAGC |
| **Primers/assays used in iPSC-derived models** | | |
| **Target / Section** | **Primer** | **Sequence (5'-3')** |
| *UNC13A* RT-PCR | Forward | GACATCAAATCCCGCGTGAA |
| *UNC13A* RT-PCR | Reverse | CATTGATGTTGGCGAGCAGG |
| *STMN2* RT-PCR | Forward | GCTCTCTCCGCTGCTGTAG |
| *STMN2* RT-PCR | Reverse | CGAGGTTCCGGGTAAAAGCA |
| *STMN2* RT-PCR | Cryptic reverse | CTGTCTCTCTCTCTCGCACA |

| | | |
|---|---|---|
| *TARDBP* RT-qPCR (SYBR Green) | Forward | GATGGTGTGACTGCAAACTTC |
| *TARDBP* RT-qPCR (SYBR Green) | Reverse | CAGCTCATCCTCAGTCATGTC |
| *GAPDH* RT-qPCR (SYBR Green) | Forward | CACCAGGGCTGCTTTTAACT |
| *GAPDH* RT-qPCR (SYBR Green) | Reverse | GACAAGCTTCCCGTTCTCAG |
| *TARDBP* RT-qPCR (Taqman; JUN) assay | - | Hs00606522_m1 (Thermo) |
| *GAPDH* RT-qPCR (Taqman; JUN) assay | - | GAPDH-Jun 4485713 (Thermo) |
| *STMN2* total RT-qPCR (exon 1-2; Taqman; FAM) assay | - | Hs00199796_m1 (Thermo) |
| *STMN2* correctly spliced RT-qPCR (exon 2-3; Taqman; VIC) assay | - | Hs00975900_m1 (Thermo) |
| *UNC13A* total RT-qPCR (exon 25-26; Taqman; FAM) assay | - | Hs00392638_m1 (Thermo) |
| *UNC13A* correctly spliced RT-qPCR (exon 20-21; Taqman; VIC) assay | - | Hs00606522_m1 (Thermo) |
| **Primers used in mouse model experiments** | | |
| **Target / Section** | **Primer** | **Sequence (5’-3’)** |
| Cryptic *Sort1* | Forward | AAATCCCAGGAGACAAATGC |
| Cryptic *Sort1* | Reverse | GAGCTGGATTCTGGGACAAG |
| Alt. *Tsn* | Forward | GGTCTTCCTGGCAGCATTTG |
| Alt. *Tsn* | Reverse | TTGACAGACAGCCTCGATGC |
| *Tardbp* | Forward | GGGGCAATCTGGTATATGTTG |
| *Tardbp* | Reverse | TGGACTGCTCTTTTCACTTTCA |
| Cryptic *Stmn2* | Forward | GCCTTACTCAGACTCCTCTCTC |
| Cryptic *Stmn2* | Reverse | TCTTCTGCCGAGTCCCATT |
| Canonical *Stmn2* | Forward | GCAATGGCCTACAAGGAAAA |
| Canonical *Stmn2* | Reverse | GGTGGCTTCAAGATCAGCTC |
| *Gapdh* | Forward | CATGGCCTTCCGTGTTCCTA |
| *Gapdh* | Reverse | CCTGCTTCACCACCTTCTTGAT |
| *Rplp0* | Forward | ACTGGTCTAGGACCCGAGAAG |
| *Rplp0* | Reverse | CTCCCACCTTGTCTCCAGTC |
| **Primers for genotyping** | | |
| **Target / Section** | **Primer** | **Sequence (5’-3’)** |
| *hStmn2* | Forward | TGGATTACAGAATATTTCACTTCCAA |
| *hStmn2* -WT | Reverse | CCCACCCACACACATATTCAC |
| *hStmn2* -Mutant | Reverse | TTGTCAACTGTGCCACAAGC |
